# Differences in sensitivity to EGFR inhibitors could be explained by described biochemical differences between oncogenic Ras mutants

**DOI:** 10.1101/005397

**Authors:** Edward C. Stites

## Abstract

Emerging data suggest different activating Ras mutants may have different biological behaviors. The most striking example may be in colon cancer, where activating KRAS mutations generally indicate a lack of benefit to treatment with EGFR inhibitors, although the activating KRAS G13D mutation appears to be an exception. As KRAS G13D generally shares the same biochemical defects as the other oncogenic KRAS mutants, a mechanism for differential sensitivity is not apparent. Here, a previously developed mathematical model of Ras mutant signaling is used to investigate this problem. The purpose of the analysis is to determine whether differential response is consistent with known mechanisms of Ras signaling, and to determine if any known features of Ras mutants provide an explanation for differential sensitivity. Computational analysis of the mathematical model finds that differential response to upstream inhibition between cancers with different Ras mutants is indeed consistent with known mechanisms of Ras biology. Moreover, model analysis demonstrates that the subtle biochemical differences between G13D and G12D (and G12V) mutants are sufficient to enable differential response to upstream inhibition. Simulations suggest that wild-type Ras within the G13D mutant context is more effectively inhibited by upstream inhibitors than when it is in the G12D or G12V contexts. This difference is a consequence of an elevated K_m_ for the G13D mutant. The identification of a single parameter that influences sensitivity is significant in that it suggests an approach to evaluate other, less common, Ras mutations for their anticipated response to upstream inhibition.

## Introduction

Inhibition of the EGFR signaling pathway with EGFR targeting antibodies, such as cetuximab, has been shown to benefit patients with colorectal cancer [1]. EGFR is a receptor tyrosine kinase, and activation of the EGFR kinase domain, such as would occur with ligand binding or (potentially) by an EGFR mutation, results in the activation of multiple signaling pathways. Among these is the RAS/RAF/MEK/ERK Mitogen Activated Protein Kinase (MAPK) cascade. Activation of this pathway is believed to promote cell-cycle entry, and aberrant, constitutive activation of this pathway is believed to be essential to the development of cancer [2]. Compounds that reduce activation of this pathway have been developed as anti-cancer agents. Many, like cetuximab, are now in clinical use.

Resistance to targeted therapies, either *de novo* or acquired, is a major obstacle to the achievement of longstanding benefit from targeted molecular therapies. Several predictors of resistance have been identified, and many of these predictors appear logical with respect to a general understanding of the targeted signal transduction pathway. For example, constitutively active, oncogenic, KRAS mutants have been shown to indicate resistance to cetuximab in colon cancer [3]. As KRAS is downstream of EGFR in the EGFR to KRAS to MAPK cascade, this appears sensible because the oncogenic KRAS mutants are intrinsically capable of activating the same MAPK pathway that EGFR activation would induce in patients with only wild-type KRAS (WT KRAS) (Figure 1A). Indeed, KRAS mutations have been shown to indicate resistance to EGFR inhibitors in other cancers, such as lung cancer [4].

**Figure 1.**
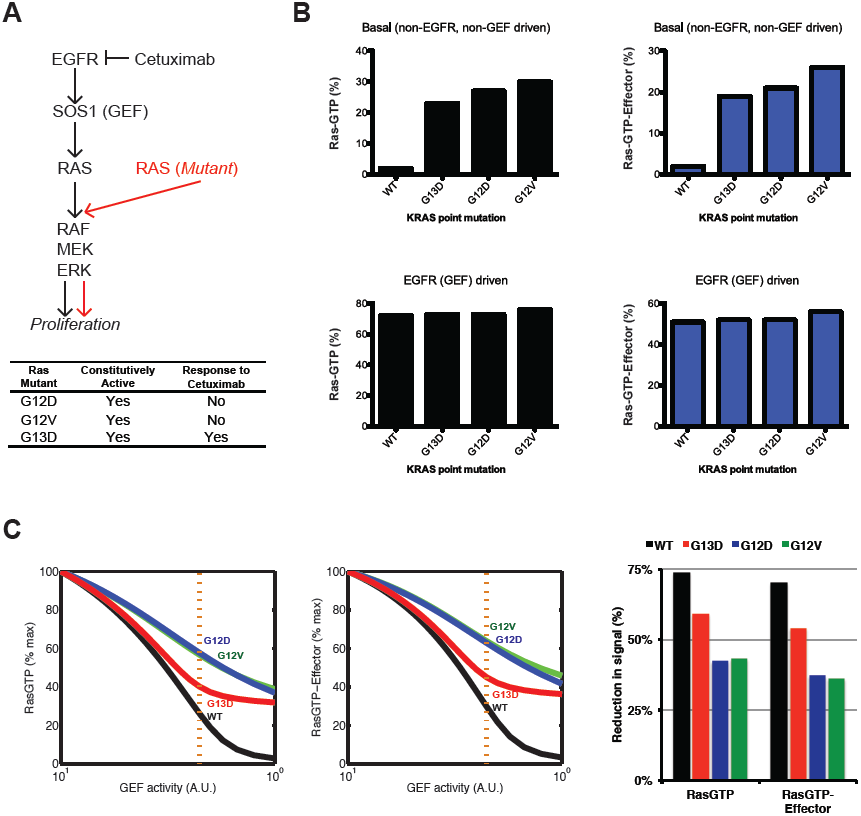
The mathematical model of Ras signaling reproduces the differential response of Ras mutants to upstream inhibitors. A) The EGFR signaling pathway and sensitivity to treatment by the EGFR inhibitor, cetuximab, for the three most common Ras mutants. B) Ras network activation due to Ras mutants G13D, G12D, or G12V as well as the all wild-type (WT) condition. (above) Basal, non-stimulated conditions. The percent of total Ras bound to GTP (left) and the percent of total effector bound to Ras-GTP (right) are both displayed as measures of Ras network activation. (below) Same as above, but for EGFR (GEF) driven conditions. All values are found with simulations of the Ras network mathematical model. C) Simulated dose responses for inhibition of the Ras network by upstream (GEF) inhibition. Networks with Ras G13D, G12D, G12V and all Ras wild-type were considered (left). Levels of RasGTP and RasGTP-effector are normalized to the maximum value. Levels of reduction from maximum to the degree of inhibition indicated with the orange, vertical, dotted line are plotted (right).

Several recent studies reveal that the relationship between oncogenic KRAS mutants and the response to cetuximab is more complicated [5–9]. Originally, a retrospective analysis performed by De Roock *et al* found that cetuximab appeared to benefit colorectal patients that had a KRAS G13D mutation [5] (Figure 1A). KRAS G13D is the third most common oncogenic KRAS mutation found in colorectal cancer. The finding that these patients had benefited from cetuximab was quite surprising, as the G13D Ras mutants are constitutively active, just like the first and second most common Ras mutants in colon cancer, G12D and G12V [10]. Interestingly, an accompanying xenograft model that employed isogenic cells in which G13D or G12D had been introduced into the endogenous locus similarly displayed differential sensitivity. However, without a logical biochemical mechanism to explain the differential response it remained unclear if the results reflected actual biology. The veracity of the initial study was later substantiated with an additional retrospective analysis of a different clinical trial dataset of colorectal cancer patients treated with cetuximab; this analysis performed by Tejpar *et al* also found that KRAS G13D patients benefit from cetuximab [6]. Several studies in tissue culture have similarly revealed a greater response to cetuximab for colon cancer cell lines that contain the KRAS G13D mutants compared to colon cancer cell lines that contain other KRAS mutants [7, 8].

The lack of a mechanism to explain the different responses of these different mutants has limited the extent to which this data can impact cancer treatment. Although empirical evidence for differential response to cetuximab has increased following the initial report by De Roock and colleagues, a mechanistic understanding of how differential sensitivity results is lacking. Indeed, a recent review by leading Ras biologists representing the National Cancer Institute’s new RAS Program declares that the differential response “challenges our understanding of how these Ras mutations actually function in clinical situations” [11]. An editorial accompanying the retrospective analysis of Tejpar *et al* opined that a biological explanation is needed before a biomarker can be used in clinical practice [12]. It is widely stated that clinical trials are needed before the retrospective analyses should influence patient management [5, 6, 9, 12, 13]. Such an opinion appears consistent with the best practices of evidence-based medicine. However, many recommended medical practices in oncology were not tested in prospective, randomized, controlled, clinical trials [14]; and studies have further found such evidence to be rare for oncology patients being treated after relapse [15]. The apparent higher standard needed for KRAS G13D likely follows from the lack of a biological basis for differential sensitivity to cetuximab.

Mathematical models that simulate the biological system have been used to study complicated biological processes and generate insights regarding a wide variety of problems in cancer biology, for example [16–23]. Mathematical models of cell signaling networks have been used to study how perturbations associated with cancer (including mutations and changes in expression) alter cell signaling, for example [24–32]. The Ras network, specifically, has been modeled as a part of larger models of growth factor signaling networks, for example [24, 33, 34]. Models of Ras signaling more specifically have provided a solid theoretical understanding for many processes in Ras biology, ranging from how membrane localization affects Ras activation [35], to Ras mediated positive feedback on its direct upstream regulator [36], to the function of Ras signaling from membrane micro-domains [37], to the cycling of Ras between different intracellular membranes [38].

It has previously been shown that a computational model that takes into account the different processes that influence the Ras nucleotide binding state can be used to uncover novel insights regarding oncogenic Ras signaling [26]. Previous application of the model focused on Ras mutants G12V and G12D. This mathematical model was revisited to evaluate whether it could contribute to the effort to explain the differential response displayed by the G13D Ras mutant. Although the G13D mutant has not been as well characterized as G12D and G12V, several properties have been at least partially characterized. Model analysis found that the described differences were sufficient to indicate differential sensitivity. Further analysis uncovered the key, measurable, biochemical property that results in differential sensitivity. Additionally, the analysis identified two other Ras mutants that have been described in colon cancer that have similar properties and should be considered for differential response. Several testable hypotheses and experiments for further evaluation are also presented.

## Methods

### Mathematical Model and Analysis

Details of the model and its development have been published previously [26, 39-41]. Briefly, the model focuses on Ras and the types of proteins that directly interact with Ras to regulate Ras GTP levels: Ras GEFs (e.g. Sos1), Ras GAPs (e.g. neurofibromin/NF1), and Ras effector proteins (e.g. RAF). The model includes 1) GEF mediated nucleotide exchange, 2) intrinsic nucleotide exchange, 3) GAP mediated nucleotide hydrolysis, 4) intrinsic nucleotide hydrolysis, and 5) effector binding. Reaction mechanisms modeled are presented below with numbering of the reaction corresponding to the process indicated above:

1. RasGDP + GEF <−> RasGTP + GEF
2. RasGXP <−> Ras()
3. RasGTP + GAP −> RasGDP + GAP
4. RasGTP −> RasGDP
5. RasGTP + Effector <−> RasGTP-effector

GXP could indicate GTP or GDP for 2, above. Free GTP, GDP, and P_i_ are not indicated in the biochemical reactions above for simplicity and are assumed to be present at constant concentrations. GEF and GAP reactions, 1 and 3 above, are described mathematically with competitive Michaelis-Menten kinetics of the reversible and irreversible types, respectively. The other reactions, 2,4, and 5 above, are described with first- and/or second-order mass-action kinetics. It is assumed that wild-type and Ras mutant proteins have identical reaction mechanisms as indicated above, and that differences in rate constants (or enzymatic parameters) for the reactions account for described differences. For example, Ras mutant protein G12V does not experience an increase in GTP hydrolysis in the presence of GAP despite having an appreciable affinity for GAP. In this case, the K_m_ is as measured, and k_cat_ used is the rate constant for spontaneous (GAP independent) nucleotide hydrolysis so that binding to GAP results in no change to the rate in the absence of GAP. All reactions are grouped as a set of differential equations and the amount of RasGTP (or RasGTP effector) is solved for a given set of conditions.

Parameters of the model for proteins correspond to biochemically observable properties. Approximate concentrations for the total amount of Ras GTPase, effectors, and basally active GEF and GAP have previously been used and published [41]. Rate constants and enzymatic properties (e.g. K_m_) for wild-type Ras proteins have been previously used and published [41]. Mutant proteins can be characterized by their difference from wild-type proteins in terms of a multiplicative factor, α. Values for α are determined from previous experimental studies that measured the desired property for both wild-type and mutant Ras proteins [42, 43]. For G12V and G12D, the α values that were previously determined were again used here [26]. Codon 61 Ras mutants were characterized previously [44], and these mutants are modeled with α obtained from these reported measurements (Table I).

**Table 1.** Parameters used to model oncogenic Ras mutants.

| | $k_{\text{GTPase}}$ | | $k_{\text{d,GDP}}$ | | $k_{\text{d,GTP}}$ | | $k_{\text{a,GDP}}$ | | $k_{\text{a,GTP}}$ | | $K_{\text{m,GAP}}$ | | $k_{\text{cat,GAP}}$ | | $K_{\text{d}}$ | |
| --- | --- | --- | --- | --- | --- | --- | --- | --- | --- | --- | --- | --- | --- | --- | --- | --- |
| **G12V** | 0.15 | 1 | 0.31 | 1 | 0.80 | 1 | 2.27 | 1 | 4.14 | 1 |  |  | *n.o.e.* | 2 | 0.44 | 3 |
| **G12D** | 0.40 | 1 | 0.48 | 1 | 5.00 | 1 | 1.37 | 1 | 3.43 | 1 |  |  | *n.o.e.* | 2 |  |  |
| **G13D** | 0.40 | ** | 3.00 | 4 | 3.00 | 4 |  |  |  |  | 100 | 5 | *n.o.e.* | 2 |  |  |
| **Q61H** | 0.09 | 6 | 1.48 | 6 | 0.61 | 6 |  |  |  |  | 0.71 | 6 | *n.o.e.* | 2 |  |  |
| **Q61K** | 0.02 | 6 | 1.40 | 6 | 0.60 | 6 |  |  |  |  | 57.14 | 6 | *n.o.e.* | 2 |  |  |
| **Q61L** | 0.02 | 6 | 2.46 | 6 | 1.07 | 6 |  |  |  |  | 0.06 | 6 | *n.o.e.* | 2 | 0.78 | 3 |
| **Q61P** | 0.08 | 6 | 1.05 | 6 | 1.27 | 6 |  |  |  |  | 1.43 | 6 | *n.o.e.* | 2 |  |  |
| **Q61R** | 0.02 | 6 | 1.14 | 6 | 0.45 | 6 |  |  |  |  | 42.86 | 6 | *n.o.e.* | 2 |  |  |
| **Q61W** | 0.07 | 6 | 1.00 | 6 | 0.80 | 6 |  |  |  |  | 2.86 | 6 | *n.o.e.* | 2 |  |  |

The factors presented in the table (α) indicate the multiple to which the wild-type parameter is altered. References indicate the studies in which these values are derived. If no evidence of change is available, no change from WT (α=1) is assumed (areas shaded in gray).
*n.o.e*. no observable effect. Ras GAPs are unable to catalyze GTP hydrolysis for Ras mutants at codon 12, 13, and 61^7^ for a Ras/GAP complex, k_GTPase_ was used for k_cat,GTP_ so that there was no increase due to GAP binding, and also so that there was no decrease in intrinsic GTPase activity due to GAP binding.
1. J. F. Eccleston, K. J. Moore, G. G. Brownbridge et al., *Biochem Soc Trans* **19** (2), 432 (1991).
2. E. C. Stites, P. C. Trampont, Z. Ma et al., *Science* **318** (5849), 463 (2007).
3. E. Chuang, D. Barnard, L. Hettich et al., *Mol Cell Biol* **14** (8), 5318 (1994).
4. A. Palmioli, E. Sacco, C. Airoldi et al., *Biochem Biophys Res Commun* **386** (4), 593 (2009).
5. L. Gremer, B. Gilsbach, M. R. Ahmadian et al., *Biol Chem* **389** (9), 1163 (2008).
6. S. Donovan, K. M. Shannon, and G. Bollag, *Biochim Biophys Acta* **1602** (1), 23 (2002).
7. A. G. Stephen, D. Esposito, R. K. Bagni et al., *Cancer Cell* **25** (3), 272 (2014).

Parameters for Ras G13D had to be estimated for this current study. Previous experiments described Ras G13D to have an approximately three-fold elevated nucleotide dissociation rate compared to wild-type Ras [45]. Previous studies have also described Ras G13D to be insensitivity to Ras GAP [10], and to have no appreciable binding to the Ras GAP NF1 [46]. A 100-fold increase in the K_m_ of GAP on Ras G13D is used to model the immeasurable binding to the Ras GAP NF1 (∼50-fold changes in other Ras mutants have previously been measured [44], so it is assumed that the difference must be even larger). The decreased GTPase activity of the G12D mutant is used for the G13D mutant because no specific α factor appears to have been previously measured; using the same value also better enables focus on the known biochemical differences.

Computational “chimeric” mutants are modeled mutants that have properties of two (or more) distinct Ras mutants. For example, a chimeric Ras mutant may be modeled with all of the properties of Ras G12D, except for the faster intrinsic nucleotide dissociation properties of G13D. Such a chimera could be used to evaluate how faster nucleotide dissociation would influence signaling.

The Ras network within the colorectal cancer context is assumed to be EGFR driven, and EGFR is assumed to activate Ras via increased activation of Ras GEFs like Sos1. We use a ten-fold increase in V_max_ for GEF reactions to indicate EGFR activation, as done previously to model receptor tyrosine kinase mediated Ras activation [26]. To simulate an EGFR inhibition dose response, levels of GEF activity between the ten-fold (10×) increase and the basal (1×) level are considered and the resulting level of RasGTP determined via model simulation.

RasGTP and RasGTP-effector complex are considered as measures of Ras pathway activation. Model simulations are used to determine levels of RasGTP and RasGTP-effector. Simulations and analysis are performed in MATLAB (7.11.0.584, MathWorks). The model is simulated to identify steady-state levels of Ras signaling for the specified set of network proteins.

We assume that the three Ras proteins, HRAS, NRAS, and KRAS, share similar biochemistry and can be modeled with the same set of biochemical properties; such an assumption is consistent with measurements of the three Ras proteins [47, 48]. We assume that measurements for the biochemical differences of a Ras mutant apply to that same point mutant for all three Ras genes. We assume that more than one Ras gene is expressed in colorectal cancer cells. This is consistent with many data [49], including that KRAS and NRAS mutants have been observed in colorectal cancer [50]. We here model Ras mutants as being heterozygous, such that for a KRAS mutant, one half of total KRAS will be mutant and one half of total KRAS will be wild-type. Here, we assume that 50% of total Ras is KRAS (and that 25% of total Ras is mutant).

Of note, many processes are not considered with the model. Just as studies of cancer biology using mouse models must be evaluated for how the mouse context may influence results and extrapolation to the human condition, it is important to consider how processes not included in the model influence results. For this study, the goal was to determine if biochemical differences known for Ras mutants could be sufficient to determine differential response to upstream signaling. Analysis was undertaken with the belief that the differential response of KRAS G13D appears to defy known biology, and that it would be a significant finding if known biology was indeed consistent with a differential response.

## Results

### Ras G13D mutant displays elevated Ras pathway activation similar to Ras G12V and Ras G12D

The model was simulated to study how Ras mutants G13D, G12D, and G12V affected basal, steady state Ras signaling compared to networks with only Ras wild-type (WT). Mutants were described with the parameters in Table I. Levels of RasGTP and levels of RasGTP-effector complex were both considered as measures of Ras pathway activation. In basal conditions, we find that previously described biochemical changes for RasG13D are sufficient to result in its constitutive activation at a level similar to RasG12V and RasG12D (Figure 1B, **above**).

The consequences of RasG13D, RasG12D, and RasG12V mutations were also considered in the case of a Ras network within a cancer cell with elevated EGFR activation. Elevated EGFR activation appears to be a characteristic of colon cancer, as supported by its response to EGFR inhibitors. As activation of the EGFR results in recruitment of Ras GEF Sos1 to the cytoplasmic membrane, where it can then activate Ras [35] we can model the elevated EGFR pathway activation by increasing the level of GEF (e.g. Sos1) activity in our model [26]. Within modeled EGFR/GEF driven conditions, we find that the three mutant containing networks, as well as the WT Ras network, all display similar levels of elevated Ras signal (Figure 1B**, below**).

### Simulations find Ras G13D is more sensitive than Ras G12D and Ras G12V to upstream inhibition

Dose response curves to upstream inhibition were obtained by simulating the mathematical model of the Ras network for different levels of upstream (GEF) activation remaining after inhibition (Figure 1C). These were performed for the Ras WT case (where all Ras was wild-type), and for mutant Ras G12D, G12V, and G13D cases. The resulting dose response curves matched well with what has been observed clinically. The dose response curve for the modeled WT Ras network demonstrate a large reduction in Ras signal as upstream activation is inhibited, consistent with the sensitivity of wild-type Ras colorectal cancer to cetuximab. The dose response curves also demonstrate a lesser reduction in total Ras signal as upstream activation is inhibited for modeled networks containing G12D or G12V Ras mutants. This is consistent with the lack of response to KRAS mutant G12D and G12V containing colorectal cancers to cetuximab.

Most interesting, the modeled network containing the G13D Ras mutant experienced a reduction in Ras signal that was greater than what was observed for mutant G12D and G12V networks, but was less than that observed for a modeled network with all WT Ras. This is consistent with previous experimental results in tissue culture systems [7, 8], with results from xenograft model systems [5], and with the different clinical responses for KRAS G13D compared to KRAS G12D and KRAS G12V containing colorectal cancers [5, 6, 13].

### Simulations reveal the elevated K_m_ for Ras GAP on Ras G13D is responsible for differential sensitivity

As the differences in the simulated dose response must follow from the differences in the parameters of the modeled mutant proteins, we simulated an extended set of “chimeric” mutants to identify which property/properties caused the differential response to upstream inhibition. For example, to evaluate whether the elevated dissociation rates of G13D contributed to the differential response, a “computational chimera” was generated by using the rate constants for G13D except for the rate constants that characterize the intrinsic nucleotide dissociation and association, which were replaced with the corresponding values of the G12V mutant (Figure 2A). (Similarly, the intrinsic nucleotide exchange properties of G13D could be combined with the other properties of the G12V or G12D mutant to also evaluate the contribution of the differences in intrinsic nucleotide exchange rates on differential sensitivity to upstream inhibition.) To study the effects of K_m_, the K_m_ of G13D could be used with the G12D and G12V mutants, or the K_m_ of wild-type Ras (which is the same for G12D and G12V) could be used for G13D. To succinctly capture the behavior of these chimeric mutants, we define “delta” (Δ) as the largest difference between the G12D mutant and the modeled chimeric mutant. A negative delta indicates a greater response to upstream inhibition.

**Figure 2.**
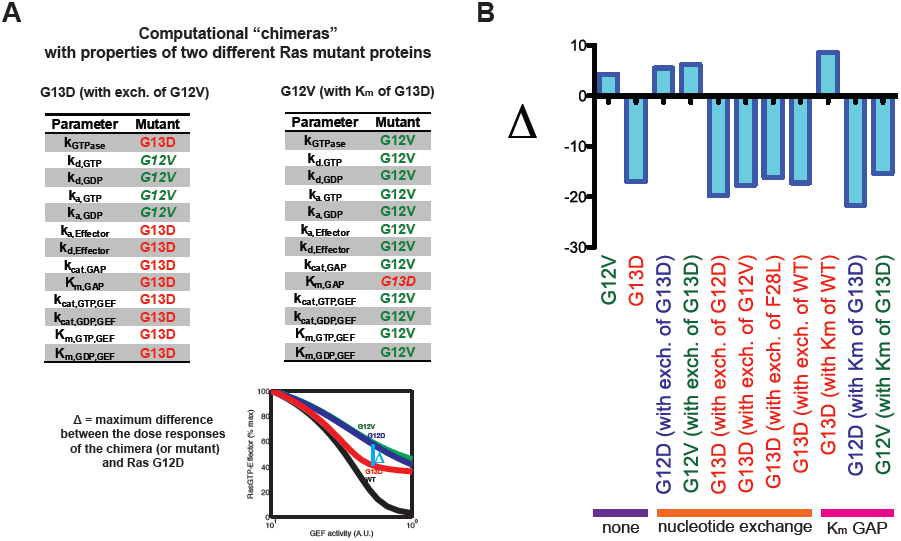
The modeled differential sensitivity follows from an elevated K_m_ for the Ras/GAP interaction. A) The contributions of different Ras signaling biochemical parameters were assessed by considering computational “chimeras” that included properties of more than one Ras mutant protein. For example, a chimera could have all of the properties of Ras G13D with the exception of having the intrinsic nucleotide exchange properties of Ras G12V (left). Or a chimera could have all of the properties of Ras G12V, except for having the Michaelis constant (K_m_) for the activity of GAP on Ras from the Ras G13D mutant. To efficiently consider multiple dose responses for different chimeras, the maximum difference between the dose responses (Δ) can be considered (bottom). B) The dose responses of considered computational chimeras find that the elevated Michaelis constant (K_m_) of Ras G13D is most responsible for the greater sensitivity to upstream inhibition, and that the elevated nucleotide exchange rate has a much smaller, and opposite, effect. Colors indicate the mutant that provided the majority of the parameters for the chimera (G12D, blue; G12V, green; G13D, red).

When nucleotide exchange was considered, modeling revealed that the elevated intrinsic nucleotide exchange of G13D did not contribute to its increased sensitivity to upstream inhibition (Figure 2B). On the contrary, modeling revealed that elevated intrinsic nucleotide exchange resulted in a lessened response to upstream inhibition. For example, replacement of the G13D parameters with the exchange parameters of G12D or G12V resulted in a chimeric mutant that was modestly more responsive to upstream inhibition. Similarly, replacement of G12D (or G12V) parameters with the exchange parameters of G13D made those mutants even less responsive to upstream inhibition.

In contrast, the chimeras provided evidence that the Michaelis constant (K_m_) for the Ras G13D interaction with Ras GAP is responsible for the increased response to upstream inhibition (Figure 2B). Replacement of this value in the G13D mutant with the value of wild-type Ras (which is the same as the value used for G12V and G12D mutants) found this computational chimera was no longer more sensitive to upstream inhibition, and was actually less responsive to upstream inhibition that Ras G12D. Modeling the chimeras formed by the replacement of the K_m_ of Ras G12D and Ras G12V with the K_m_ of Ras G13D found these two chimeric mutants to be more sensitive to upstream inhibition.

### A differential ability to activate wild-type Ras underlies the differential sensitivities of G13D vs. G12D and G12V Ras to cetuximab

The original dose-response of G13D, G12D, and G12V was further evaluated to determine how an elevated K_m_ might result in differential sensitivity. As a mutant-containing cell, as well as the mutant-containing network, contains both mutant and wild-type Ras, we considered the dose response for both the mutant and wild-type fractions of Ras. Intriguingly, the simulations found that the mutant fraction of Ras was not significantly affected by upstream inhibition for mutants G13D, G12D, or G12V (Figure 3A, **above**). This is consistent with intuition, where it does not seem that upstream inhibition should influence Ras mutant signaling. However, wild-type Ras signaling was largely affected by upstream inhibition (Figure 3A**, below**). The G13D mutant network displayed a more rapid decline in wild-type Ras signal levels than did the fully wild-type network or for the Ras G12D and RasG12V networks. Of note, the G13D network displayed a near complete reduction of wild-type RasGTP-effector signal, like the wild-type network, whereas the G12D and G12V networks maintained a modestly elevated level of wild-type RasGTP-effector signaling at full upstream-induced GEF inhibition (Figure 3A**, below**).

**Figure 3.**
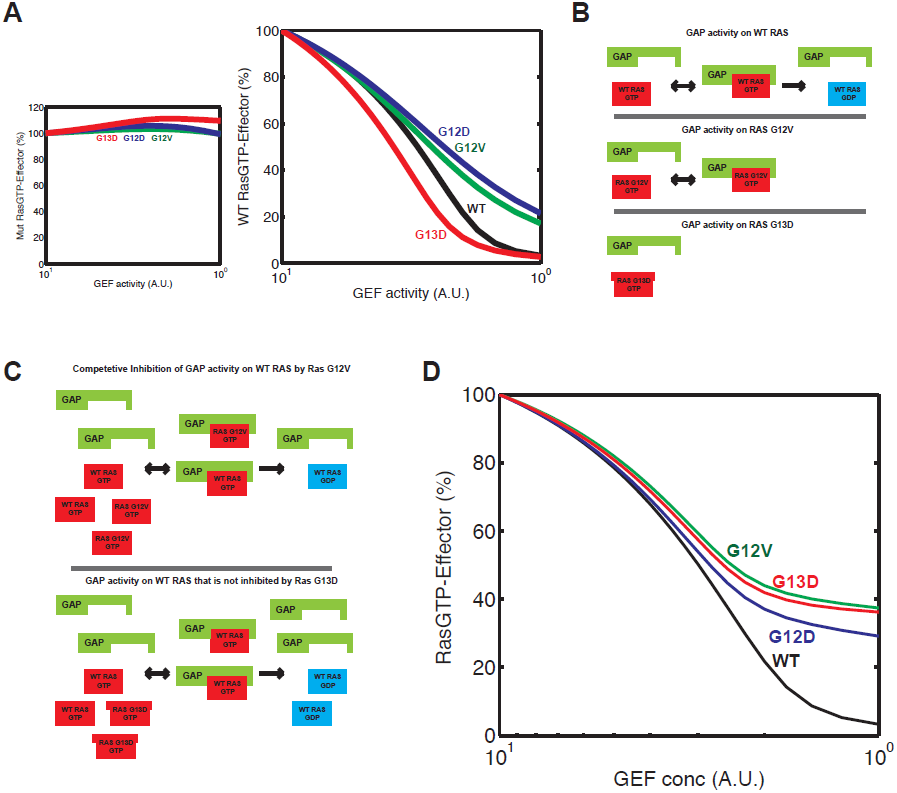
Modeling suggests differential sensitivity follows from differences in wild-type Ras signaling. A) Levels of Ras pathway activation for mutant and wild-type subsets of total Ras for an upstream inhibition dose response. The Ras network is indicated by the mutant present (or WT if no mutant present). (above) Ras mutants experience minimal changes in activity level as a consequence of upstream inhibition. Wild-type Ras in the different contexts display large decreases in the active fraction (below). The wild-type Ras in the G13D mutant containing network displays the earliest response during the simulated dose response. B) Differences in the Ras GAP interaction with wild-type (WT), mutant G12V, and mutant G13D Ras. (above) Ras GAPs bind to WT RasGTP, form a GAP-Ras complex, and facilitate GTP hydrolysis. (middle) GAP insensitive Ras mutants, like Ras G12V, are typically capable of interacting with Ras GAP, but Ras GAP has no significant effect on GTP hydrolysis. (below) Ras G13D has been reported to have a significantly reduced interaction with the Ras GAP neurofibromin (NF1). C) Competitive inhibition of Ras GAPs as a mechanism that increases WT RasGTP. Constitutively active Ras mutants that bind to Ras GAPs sufficiently well can form transient, non-productive, GAP-Ras complexes. This effectively reduces the pool of available Ras GAPs to inactivate any WT RasGTP, and the dynamic equilibrium therefore settles into a new steady-state with higher levels of WT RasGTP. A constitutively active Ras mutant with impaired binding to Ras GAPs, such as has been reported for Ras G13D, is a much less effective competitive inhibitor of Ras GAPs and WT RasGTP experiences a negligible increase in the presence of such a mutant. D) G12D and G12V mutant containing networks respond to upstream inhibition if competitive inhibition is removed from the model.

In our previous analysis of oncogenic Ras mutant signaling, we proposed an additional mechanism by which oncogenic Ras mutants contribute to the elevated levels of Ras signaling: the competitive inhibition of Ras GAPs by the GTP-bound Ras mutant leading to increased wild-type RasGTP [26]. At basal, unstimulated, conditions, Ras GAPs catalyze the conversion of RasGTP to RasGDP to maintain a low level of Ras GTP (Figure 3B). Constitutively active Ras mutants at codon 12, 13, or 61 have impaired intrinsic GTPase activity and are insensitive to GAP catalyzed GTP hydrolysis [11]. The RasGTP bound mutant, however, typically maintains the ability to bind to Ras GAPs. The transient interaction between a GAP-insensitive mutant and a Ras GAP reduces the pool of available Ras GAPs that can act on any wild-type RasGTP, shifting the dynamic equilibrium to a higher level of RasGTP (Figure 3C). Increased wild-type Ras signaling in combination with oncogenic Ras mutants has been demonstrated multiple times by multiple groups [26, 51–55]. Ras G13D may not be able to competitively inhibit GAPs to as large of an extent because it has been reported that RasG13D does not detectably bind the Ras GAP neurofibromin [46].

To confirm that competitive inhibition within the model was the process responsible for the modeled differential response, the competitive inhibition term was removed from the equations describing Ras GAP activity in the same manner it was removed in our original manuscript to evaluate the contribution of competitive inhibition to Ras G12D and Ras G12V activation [26]. Upon removal of competitive inhibition, Ras G12V was now nearly as responsive as Ras G13D to upstream inhibition, and Ras G12D was found to be even more responsive than Ras G13D to upstream inhibition (Figure 3D).

### Consideration of codon 61 mutants confirms that elevated K_m_ leads to a differential response to upstream inhibition

Many KRAS mutants other than G13D, G12D, and G12V have been observed in colorectal cancer patients [56] (Figure 4A). If patients with KRAS G13D mutations are demonstrated to benefit from cetuximab in controlled, prospective, clinical trials, it will raise the important issue of how patients with other, less common, KRAS mutants should be treated if constitutive activity is not in itself sufficient to indicate a lack of benefit to cetuximab. A clinical trial for each and every distinctive KRAS mutant is not a practical option [12]. One additional value of identifying the mechanism that explains the KRAS G13D differential response is that any KRAS mutant could be evaluated for the property that determines the ability to respond to the drug to identify which patients with which mutants may benefit.

**Figure 4.**
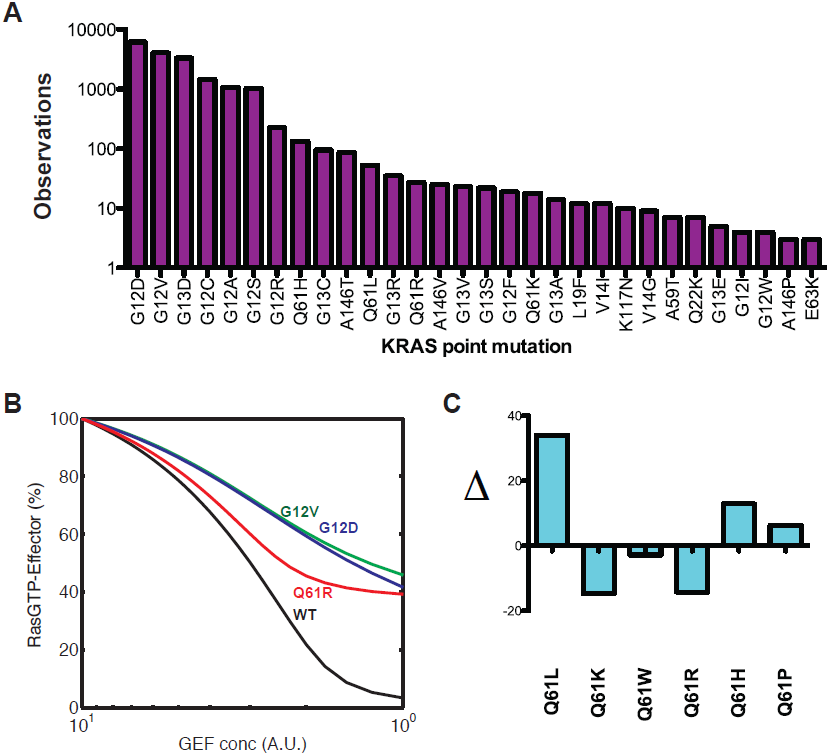
Modeling suggests other Ras mutants observed in colorectal cancer may also be sensitive to upstream inhibition. A) Incidence of different KRAS point mutations within colorectal cancer. Data from COSMIC [56] for KRAS point mutations that have been observed at least three times in colorectal cancer. G12D, G12V, and G13D are the three most common mutations that have been reported. There is a long tail of less frequently observed KRAS mutations. Of note, the frequency with which these less common mutations occur may be underestimated from this data as some of the studies in the aggregated COSMIC data may have only considered a limited number of exons, codons, and/or point mutations. B) Dose responses for codon 61 Ras mutants find the Q61R mutants, which has an elevated Michaelis constants (K_m_), displays an increased level of sensitivity to upstream inhibition relative to the G12D mutant. C) Deltas (Δ) for five characterized codon 61 Ras mutants. A negative value for Δ indicates the mutant has greater sensitivity to upstream inhibition.

Of note, six codon 61 Ras mutants have previously been characterized in terms of nucleotide exchange, GTPase activity, and GAP interaction [44]. Of these six mutants, at least four (Q61H, Q61L, Q61 K, Q61R) have been observed in colon cancer (Figure 4A). Data from these studies (Table I) were applied to our model. Consistent with the finding in the computational chimera studies that an elevated K_m_ resulted in increased sensitivity to upstream inhibition, the three mutants with the most elevated K_m_ (Q61R and Q61 K) were found to display increased sensitivity to upstream inhibition (Figure 4B,C). Additionally, the mutant with the lowest K_m_, Q61L, actually displayed the least response to upstream inhibition (Figure 4C).

## Discussion

Based upon a mathematical analysis of the reactions that regulate Ras signaling and that further incorporate the various biochemical alterations of Ras mutants, the following model is proposed to explain the different responses colorectal cancer patients have to cetuximab (Figure 5). In a colorectal cancer with all wild-type Ras, Ras signaling is driven only by EGFR activation. EGFR activation results in increased Ras GEF activity, which leads to increased RasGTP. Inhibition of EGFR activation with cetuximab reduces Ras GEF activity, and thereby reduces RasGTP, allowing for reduced pro-growth signals (Figure 5**, left**). In a colorectal cancer with a G12D or G12V mutation, the inhibition of EGFR/GEF mediated wild-type Ras activation results in only a partial reduction of wild-type Ras signal because wild-type Ras is also activated as a consequence of the competitive inhibition of Ras GAPs by the mutant Ras G12D (or G12V) (Figure 5**, center**). Within colorectal cancers with the G13D mutation, inhibition of EGFR is sufficient to reduce the RasGTP proportion of wild-type Ras to a much larger extent than in G12D or G12V containing colorectal cancers because the G13D mutant does not bind basally active Ras GAPs sufficiently well to result in an accumulation of wild-type RasGTP (Figure 5**, right**). Just as with the G12D and G12V mutants, levels of G13D activity are not significantly affected by upstream inhibition. Within this model, there are at least two possible explanations for why reduced wild-type Ras signaling causes the clinical response. One is that the total amount of Ras activation has been reduced to a level that influences cancer survival (or proliferation), which would be consistent with graded phenotypes in response to Ras activation [44, 57]. The other potential explanation is that wild-type Ras signal has distinct effects from mutant Ras, and that both signals are needed for the survival of the cancer [53, 54].

**Figure 5.**
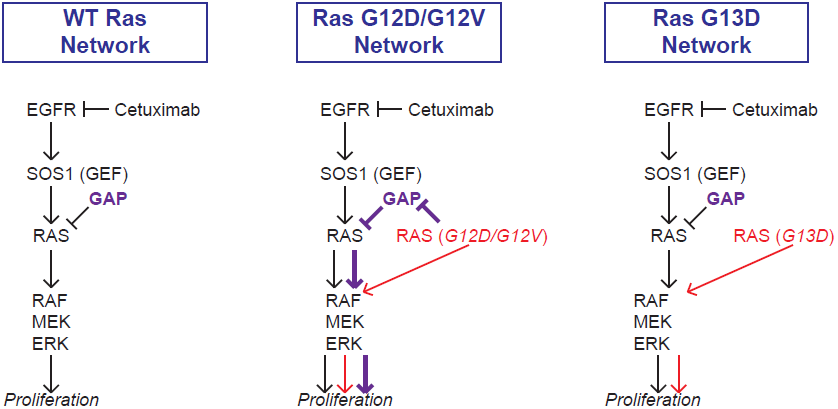
A mechanism to explain the differential sensitivity of G13D, G12D, and G12V KRAS mutants as suggested by mathematical modeling. Proposed mechanism to explain the differential sensitivities of activating KRAS mutants to cetuximab. (left) Within an EGFR driven cancer with all WT Ras, Ras activation results only as a consequence of EGFR activation. Inhibition of EGFR is sufficient to return Ras activation to low, basal, levels. (middle) Within an EGFR driven cancer that also contains an activating KRAS mutant, like G12V or G12D, there are three ways that Ras is activated: i) EGFR mediated Ras wild-type activation, ii) Ras mutant mediated activation of the mutant fraction of Ras, iii) Ras wild-type activation that follows from competitive inhibition of Ras GAPs. EGFR inhibition can only reduce i), and high levels of active mutant and wild-type Ras remain. (right) Within an EGFR driven cancer that also contains an activating KRAS that interacts minimally with Ras GAPs, as has been reported for G13D, only i) and ii) above contribute to Ras activation. EGFR inhibition can thereby eliminate WT Ras activation.

The mechanism proposed above depends on the competitive inhibition of Ras GAPs by Ras mutant G12V and Ras G12D, but not Ras G13D. As the model studied includes various simplifications, it is prudent to evaluate these simplifications and how they might influence the model predictions. Competitive inhibition is somewhat dependent upon the assumptions that basally active Ras GAPs are localized to the same membranes where Ras is signaling, and that localization to the membrane results in a higher apparent affinity of GAP for Ras than if the GAP were localized in the cytoplasm [39]. The assumption of membrane localization seems justified. For example, Spred1 has previously been shown to localize to be membrane localized [58] and Spred1 has recently been shown to mediate membrane localization of NF1 [59]. Loss of SPRED1 results in the “Rasopathy” Legius syndrome [60]; Legius syndrome has many similarities with neurofibromatosis type 1, which results from germline mutations to the Ras GAP NF1 [61]. That membrane localization increases can result in an increased apparent affinity is supported theoretically [62] and has long been believed to apply to Ras regulation [24, 35].

Competitive inhibition also depends on the total amount of basally active GAP. We assume an enzymatic level of basally active GAP. A large quantity of basally active Ras GAPs could exceed the amount of Ras mutant available to inhibit it. However, a low level of Ras GAP activity is sufficient to maintain low levels of Ras GTP in unstimulated cells with all wild-type Ras. It seems unlikely that an excess of GAP to Ras is used to maintain low levels of RasGTP in the absence of stimulus. Another simplification is that feedback is not included in the model. There are multiple positive and negative feedback loops in the EGFR to RAS to ERK pathway [63]. Feedback has been proposed to result in elevated levels of Ras GAP activity [64], and also to result in reduced levels of Ras GAP activity [65]. That elevated wild-type Ras has been identified in cell lines that contain a constitutively active mutant Ras [26, 51–55] suggests that negative feedback does not occur to a level sufficient to reduce wild-type RasGTP levels back to the levels found in cells without a Ras mutant. Although it is certainly possible that the differential sensitivity is an artifact of the model and that it is simply a coincidence that the G13D containing network has a greater sensitivity within the model, there does not appear to be a definitive relationship between a model simplification and the differential response observed.

Although the model analysis is based upon previous experimental data, it is essential that the insights and hypotheses developed with the model are subjected to experimental investigation. The central hypothesis of this work is that the differential response to cetuximab and anti-EGFR treatments between G13D colon cancers and G12D and G12V colon cancers follows from differences in wild-type activation in the presence of inhibitor. KRAS G13D containing colon cancer cell lines respond to cetuximab [7, 8]; these cells and cells with KRAS G12D or G12V could have levels of wild-type Ras GTP measured as part of a cetuximab dose response. Measurement of HRAS and NRAS GTP levels is an established approach for the measurement of wild-type Ras activation within KRAS mutant containing cells [26, 51–55].

Other experiments can test the hypothesis that the effects of cetuximab on Ras signaling are mediated primarily through Ras GEFs. A dose response with dominant negative Ras S17N mutant, which acts by inhibiting Ras GEFs [66], could evaluate if reduction of Ras GEF activity results in reduction of wild-type Ras signaling within these colon cancer cells. A greater response would be anticipated for Ras G13D containing colon cancer cells relative to Ras G12D or Ras G12V colon cancer cells.

It would be valuable to measure the K_m_ of KRAS G13D with NF1. Of note, the previous study that did not detect an interaction between the G13D mutant and GAP used HRAS G13D and NF1 [46]. Additionally, a complete characterization of KRAS G13D biochemistry would allow for identification of any, unknown, largely different biochemical properties and would enable a more complete computational study of the KRAS G13D mutant. There are many different proteins that display GAP activity on Ras [67]. It could also be valuable to measure the Ras/GAP interaction K_m_ and k_cat_ between G13D mutant and other Ras mutants for a variety of Ras GAP proteins; there appears to be a limited correlation between the K_m_ between a Ras mutant and other Ras GAPs [44]. Moreover, it is not clear which Ras GAPs besides NF1 contribute to the basal regulation of Ras GTP levels. Within prostate cancer cells, for example, DAB2IP has been shown to contribute to the regulation of basal Ras activity [68].

Finally, model-based analysis suggests KRAS Q61R and KRAS Q61 K containing colon cancer cells may respond to cetuximab like the KRAS G13D mutant. Although the Q61R and Q61 K mutants are less common than the G12D, G12V, and G13D mutants, they have been observed in colorectal cancer (Figure 4A). Whether or not patients with these mutations benefit from cetuximab could potentially be assessed with an analysis of existing clinical trial data if there are sufficient numbers of patients with these mutants that have been treated. Extension of engineered isogenic cell lines [5, 69, 70] to codon 61 Ras mutants, including the Q61R and Q61 K mutations, could provide a model system for testing these mutants in the colon cancer context. Ultimately, prospective trials could evaluate whether patients with the Q61R or Q61 K benefit from cetuximab.

The ability to identify which specific mutants are associated with a clinical response is essential to the ultimate realization of personalized cancer medicine [71–73]. Visions of personalized medicine that claim a knowledge of which genes are mutated in a patient’s cancer can be used to determine treatments for that patient are oversimplistic; not all mutations are equivalent. However, the opposite extreme, where treatment can only be determined with knowledge of each specific mutant, is problematic. For example, more than 90 different KRAS point mutations have been observed in colorectal cancer (Figure 4A). There is a long tail of infrequently seen mutations. It is not practical or feasible to conduct clinical trials for each and every mutation to determine whether or not that mutation generally indicates responsiveness.

The work presented here suggests that intermediate levels of mutant characterization could be useful. Here, an elevated K_m_ was theoretically demonstrated to result in increased sensitivity to upstream inhibitors, like cetuximab. If this hypothesis is confirmed experimentally, it presents a third line of attack for personalized medicine. The task of measuring K_m_ for all KRAS mutants that have been observed in colorectal cancer, as well as all potential hot-spot mutations, is a task that can be accomplished. KRAS mutants with an elevated K_m_ could be considered as a separate class with the hypothesis that those mutants will benefit from cetuximab. Controlled clinical trials could evaluate that hypothesis. If clinical trials demonstrate a general benefit for patients with a KRAS mutant characterized by an elevated K_m_, this information could be applied to other mutants. For example, if a novel insertion mutant were identified in a patient, the K_m_ could be measured and the results of that measurement could be considered by the clinician and patient to determine if treatment with cetuximab should be pursued.

Overall, this model-based analysis provides a demonstration for how mathematical models can be used to investigate whether or not surprising clinical and biological findings are actually consequences of known biology or rather indicate that new biology must be at play. For the specific case of the unexpected differential response of KRAS G13D to cetuximab, the analysis presented here finds that differential response is indeed consistent with known mechanisms of Ras biology, and moreover that specific, previously observed biochemical properties of KRAS G13D are sufficient to result in a differential response. Several testable hypotheses are presented. Overall, this model-based analysis provides several new leads for the continued, multi-modal, investigations into the response of cancer to targeted therapies.

